# Transmembrane helices mediate the formation of a stable ternary complex of cyt b_5_ reductase, cyt b_5_, and SCD1

**DOI:** 10.1101/2021.11.13.468331

**Authors:** Jiemin Shen, Gang Wu, Ah-Lim Tsai, Ming Zhou

## Abstract

Mammalian cytochrome b_5_ (cyt b_5_) and cytochrome b_5_ reductase (b_5_R) are electron carrier proteins required for many membrane-embedded oxidoreductases. Both cyt b_5_ and b_5_R have a cytosolic domain anchored to the membrane by a single transmembrane helix (TM). It is not clear if b_5_R, cyt b_5_ and their partner oxidoreductases assemble as binary or ternary complexes. Here we show that b_5_R and cyt b_5_ form a stable binary complex, and that b_5_R, cyt b_5_ and a membrane-embedded oxidoreductase, stearoyl-CoA desaturase 1 (SCD1) form a stable ternary complex. The formation of the complexes significantly enhances electron transfer rates, and that the single TM of cyt b_5_ and b_5_R mediated assembly of the complexes. These results reveal a novel functional role of TMs in cyt b_5_ and b_5_R and suggest that an electron transport chain composed of a stable ternary complex may be a general feature in oxidoreductases that require the participation of cyt b_5_ and b_5_R.

## Introduction

Cytochrome b_5_ (cyt b_5_) and cytochrome b_5_ reductase (b_5_R) are obligatory partners for a number of oxidoreductases such as fatty acid desaturases and elongases, oxygenases, and cytochrome P450s (cyt P450)(Schenkman and Jansson 2003, Elahian, Sepehrizadeh et al. 2014). They form part of an electron transport chain that transfers electrons from a reductant, nicotinamide dinucleotide (NADH) or nicotinamide dinucleotide phosphate (NADPH), to the flavin adenine dinucleotide (FAD) cofactor on b_5_R(Yamada, Tamada et al. 2013), the heme moiety of cyt b_5_(Vergeres and Waskell 1995) and finally the metal ions in the catalytic center of oxidoreductases.

Mammalian stearoyl CoA desaturase-1 (SCD1) is a membrane-embedded oxidoreductase that catalyzes the rate-limiting step in the formation of the first double-bond in saturated fatty acids. SCD1 has a major role in the regulation of fatty acid metabolism and membrane synthesis and is a validated drug target(Paton and Ntambi 2009). The enzymatic cycle of SCD1 involves redox transition of a diiron center, which requires participation of b_5_R and cyt b_5_ to deliver reducing equivalent from NADH (Scheme 1).

Extensive structural and functional studies have been conducted on the soluble forms of b_5_R and cyt b_5_ that lack their transmembrane (TM) segments. These studies show that the soluble domains of b_5_R and cyt b_5_ are sufficient to support electron transfer, and that charged residues on the surface of the soluble domains of b_5_R and cyt b_5_ likely mediate their interactions(Dailey and Strittmatter 1979, Strittmatter, Kittler et al. 1992, Nishida and Miki 1996, Kawano, Shirabe et al. 1998, Shirabe, Nagai et al. 1998, Samhan-Arias, Almeida et al. 2018). However, less attention has been paid to the role of their TM domains, and a stable b_5_R/cyt b_5_ binary complex has never been isolated. Studies of electron transfer from cyt b_5_ to SCD1 have been limited to demonstration that cyt b_5_ is required for the activity of SCD1(Paton and Ntambi 2009, Nagao, Murakami et al. 2019). Whether cyt b_5_ and SCD1 form a stable complex and whether a stable complex enhances the activity of SCD1 have not been explored. These questions are significant in terms of understanding how each redox component of the electron transport chain may function in the native environment and the mechanism of electron transfer. Knowledge of stable binary or ternary complexes of electron transfer partners is also relevant in developing novel strategies to inhibit membrane-bound oxidoreductases, for example, SCD1, which is a validated drug target for many types of cancers(Ackerman and Simon 2014, Theodoropoulos, Gonzales et al. 2016, Savino, Fernandes et al. 2020, Oatman, Dasgupta et al. 2021), neurodegenerative diseases(Vincent, Tardiff et al. 2018, Fanning, Haque et al. 2019, Nuber, Nam et al. 2021), and metabolic diseases(Ntambi, Miyazaki et al. 2002, Gutiérrez-Juárez, Pocai et al. 2006, Aljohani, Syed et al. 2017).

## Results

### Existence of binary and ternary complexes in cells

We first examined whether SCD1, cyt b_5,_ and b_5_R form binary or ternary complexes in cells. By fusing SCD1 with a green fluorescent protein (GFP), cyt b_5_ with a Myc tag, and b_5_R with a hemagglutinin (HA) tag, we monitored cellular localization of the three proteins by immunofluorescence confocal microscopy. When SCD1 was expressed, meshwork-like distribution of GFP fluorescence was observed (Extended Data Fig. 1), consistent with its localization to the endoplasmic reticulum (ER) membranes(Man, Miyazaki et al. 2006). Cells co-expressed with SCD1 and cyt b_5_ exhibited overlapping fluorescence, and although the fluorescence from cyt b_5_ clustered with most of that from SCD1 as shown in yellow color in the merged image (Fig 1a), cyt b_5_ seemed to have a wider distribution than SCD1 likely due to different expression levels of these proteins and participation of cyt b_5_ in multiple redox pathways. Colocalization was also evident in cells co-expressing cyt b_5_ and b_5_R (Fig. 1b), consistent with their roles in mediating electron transfer to redox enzymes. As expected, co-localization of all three proteins was also observed when all three were co-expressed in the same cells (Fig. 1c). These observations suggest that cyt b_5_ may form binary complexes with b_5_R or SCD1 and that the three may form a ternary complex.

**Fig. 1:**
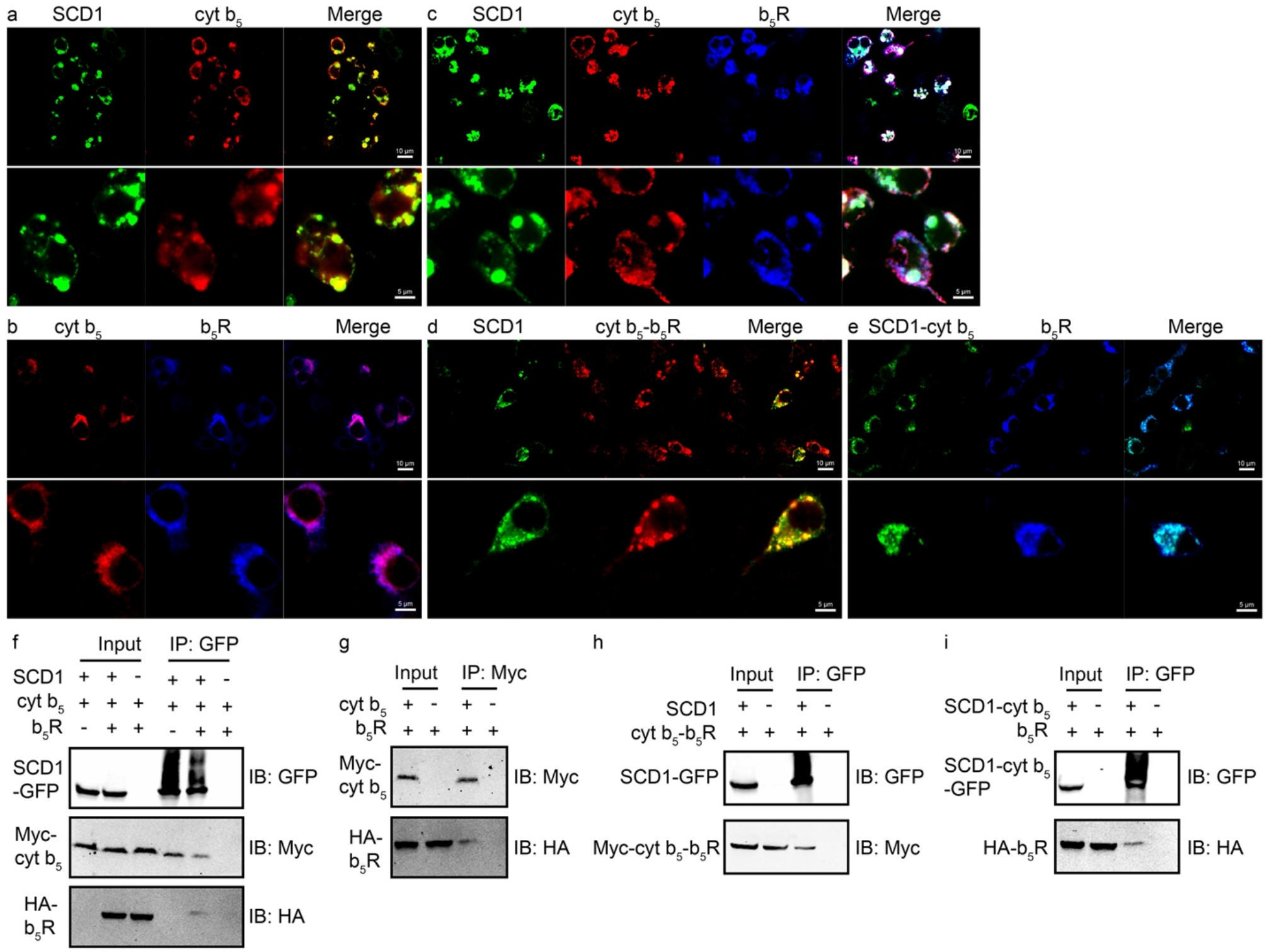
Colocalization and coimmunoprecipitation of SCD1, cyt b_5_, b_5_R, and their binary fusions. Confocal microscopy images show the subcellular distribution and colocalization of **a**, SCD1-GFP (green) and Myc-cyt b_5_ (red); **b**, Myc-cyt b_5_ (red) and HA-b_5_R (blue); **c**, SCD1-GFP (green), Myc-cyt b_5_ (red), and HA-b_5_R (blue); **d**, SCD1-GFP (green) and Myc-cyt b_5_-b_5_R (red); **e**, SCD1-cyt b_5_-GFP (green) and HA-b_5_R (blue). In **a**-**e**, images in the top and bottom panels are from same samples of different magnifications. **f**, Coimmunoprecipitation of SCD1, cyt b_5_ and b_5_R. Cells co-expressing SCD1-GFP and Myc-cyt b_5_ (left lane) were solubilized and cell lysate was immunoprecipitated with GFP nanobody resins. Detection of Myc-cyt b_5_ after extensive wash of the resins indicates some stable complex assembly between SCD1 and cyt b_5_. Similarly, weak ternary complex formation was demonstrated from cells co-expressing tagged SCD1, cyt b_5_, and b_5_R (middle lanes). Cell lysate from cells expressing tagged cyt b_5_ and b_5_R (right lane) was served as a negative control to exclude the possibility of non-specific binding of cyt b_5_ and b_5_R to resins and non-specificity of antibodies used in western blots. **g**, Coimmunoprecipitation of cyt b_5_ and b_5_R shows the existence of some stable cyt b_5_-b_5_R complex. Unlike in **f**, cell lysates were immunoprecipitated with anti-Myc antibodies and protein A resins to capture Myc-cyt b_5_. **h**, Coimmunoprecipitation of SCD1 with the binary fusion of cyt b_5_-b_5_R. **i**, Coimmunoprecipitation of the binary fusion of SCD1-cyt b_5_ and b_5_R. For all the input lanes, 6% of cell lysate was loaded. IB, immunoblotting; IP, immunoprecipitation. All the data are from one representative experiment of at least two independent repeats.

To test whether the proximity in their expression pattern leads to the formation of stable binary or ternary complexes, we next examined their interactions by co-immunoprecipitation (co-IP). We found co-IP of cyt b_5_ and SCD1, cyt b_5_ and b_5_R, and all three when either two or three proteins were co-expressed. (Fig. 1f and 1g). We also generated binary fusions by connecting SCD1 and cyt b_5_, and cyt b_5_ and b_5_R to test their assembly with b_5_R and SCD1, respectively. The SCD1-cyt b_5_ had a C-terminal GFP tag, and cyt b_5_-b_5_R an N-terminal Myc tag. Colocalization analysis showed that SCD1 and cyt b_5_-b_5_R (Fig. 1d), SCD1-cyt b_5_ and b_5_R (Fig. 1e) were in close proximity in cells. Co-IP results (Fig. 1h and 1i) indicate that the binary fusions afford stronger complex formation compared to individual SCD1, cyt b_5,_ and b_5_R.

### Stable binary complex between b_5_R and cyt b_5_

We proceeded to large-scale production of cyt b_5_ and b_5_R complex for further biochemical characterizations. However, simply co-expressing full-length cyt b_5_ and b_5_R did not produce sufficient amount of complex. To increase the yield of the complex, and encouraged by previous reports of production of stable dimeric membrane proteins after fusion of two monomers (Steiner-Mordoch, Soskine et al. 2008, Nasie, Steiner-Mordoch et al. 2010, Stockbridge, Robertson et al. 2013), we adopted the strategy of expressing a fusion protein of full-length cyt b_5_ and b_5_R as a concatenated chimera with a linker connecting the C-terminus of cyt b_5_ with the N-terminus of b_5_R (Fig. 2a). The linker contained a tobacco etch virus (TEV) protease recognition site and can be cleaved after purification. The fusion protein was expressed and purified, and the amount was sufficient for further biochemical studies (Fig. 2b left). The fusion protein contained both FAD and heme, as indicated by the UV/Vis absorption spectrum (Fig. 2b right), and eluted as a single peak on a size-exclusion column (SEC). When the linker was cleaved by TEV protease, cyt b_5_ and b_5_R stayed together as a stable complex with the same elution volume as the fusion protein (Fig. 2b left). However, when the soluble domains of cyt b_5_ and b_5_R were expressed as a fusion protein, the two soluble domains did not stay as a complex after cleavage of the linker (Extended Data Fig. 3a and 3d). These results indicate that interactions between the TM segments are required to maintain the binary complex.

**Fig. 2:**
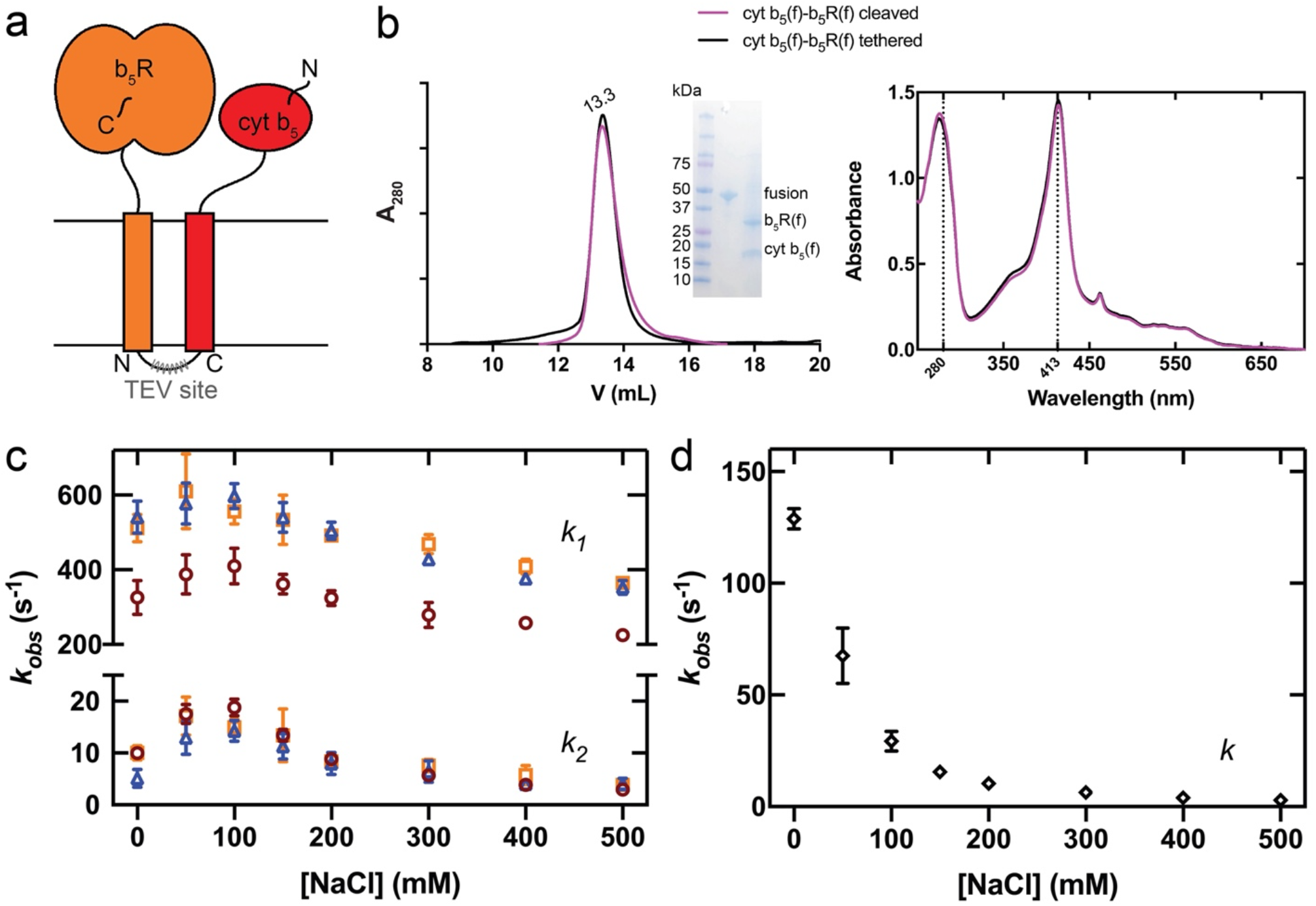
Full-length cyt b_5_ and full-length b_5_R can form a stable complex with faster electron transfer kinetics. **a**, Schematic diagram of cyt b_5_-b_5_R fusion constructs. The N and C denotes the N-terminus and C-terminus of cyt b_5_ and b_5_R domain. The gray zigzag line indicates the placement of a TEV protease site on the linker region connecting the cyt b_5_ and b_5_R. **b**, Size-exclusion chromatography (SEC) profile (left), SDS-PAGE image (inset), and UV-Vis spectra (right) of the linker-cleaved (magenta) and tethered (black) fusion of SCD1-cyt b_5_. Almost identical SEC profile and optical spectra suggest the stable assembly between cyt b_5_ and b_5_R. **c**, Ionic strength-dependent electron transfer in different constructs of cyt b_5_-b_5_R fusions: linker-cleaved full-length (orange); tethered full-length (blue); and soluble without TM domains (red). The upper and lower half of the plot represents the rate constants of the fast phase (*k*_*1*_) and slow phase (*k*_*2*_), respectively. **d**, Ionic strength-dependent electron-transfer rate between individual b_5_R and cyt b_5_. Different from those of cyt b_5_-b_5_R fusions, the time courses with individual b_5_R and cyt b_5_ are monophasic. Error bars represent standard error of the mean (SEM) from three independent repeats.

We next measured the rate of electron transfer in the stable binary complex of cyt b_5_ and b_5_R. We measured the reduction of cyt b_5_ in the context of a stable complex with b_5_R or as an individual protein mixed with b_5_R. We found that the time course of cyt b_5_ reduction can be fit with a biphasic exponential function (*k*_*1*_ and *k*_*2*_) when cyt b_5_ and b_5_R were in a binary complex (Extended Data Fig. 2a). The biphasic kinetics of electron transfer was also observed in soluble fusion of cyt b_5_ and b_5_R with similar rates with those of full-length fusion. In contrast, the time course can be fit with a single exponential function when the soluble forms of cyt b_5_ and b_5_R were added (Extended Data Fig. 2b). The observed electron transfer rate (*k*_*1*_) was ∼34-fold faster in the binary complex than that in the mixture of individual proteins in 150 mM NaCl. Thus, the formation of a stable cyt b_5_ and b_5_R complex enhances spatial proximity and the precise alignment of the two soluble domains to facilitate electron transport.

The rate of electron transfer between soluble forms of b_5_R and cyt b_5_ is known to be sensitive to ionic strength(Meyer, Shirabe et al. 1995). We next tested how the ionic strength affects the rate of electron transfer from b_5_R to cyt b_5_ in the stable complex. The time courses of cyt b_5_ reduction in the binary complex and the mixture of individual proteins were followed in buffer with different NaCl concentrations, and the rates were calculated from either double exponential fitting for the fusion proteins or single exponential fitting for the individual proteins. The full-length and soluble fusions displayed similar trend of ionic strength dependence where the electron transfer rates (*k*_*1*_ and *k*_*2*_) peaked at ∼50 mM NaCl and decreased as [NaCl] increased (Fig. 2c). The difference between no NaCl and 150 mM NaCl is only 1.04-fold. However, when the two proteins were not assembled as a complex, ionic strength has a more significant effect on the rate of electron transfer (Fig. 2d): an ∼8-fold decrease was observed from no NaCl to 150 mM NaCl. These results suggest that formation of a stable binary complex aligns the soluble domains in position for electron transfer so that electrostatic interactions have a much smaller role in guiding and facilitating the proper interactions of the soluble domains.

### Stable binary complex between cyt b_5_ and SCD1

We next investigated whether the full-length cyt b_5_ can form a stable complex with SCD1. We found that simply co-expression of the two proteins do not produce high level of SCD1-cyt b_5_ binary complex. We then applied the fusion protein strategy and linked the C-terminus of SCD1 to the N-terminus of the full-length cyt b_5_ with a TEV recognition site in the linker (Fig. 3a). The SCD1-cyt b_5_ fusion protein had sufficient yield and eluted as a single peak on a size-exclusion column (Fig. 3b and Methods). After cleavage of the linker, SCD1 and cyt b_5_ stayed together in a stable complex as indicated by the single peak on the size exclusion column; the UV-Vis spectrum of the fusion protein did not show any change after TEV protease cleavage (Fig. 3b). When the soluble domain of cyt b_5_ was fused to SCD1, the two did not stay together as a complex after the linker was cleaved (Extended Data Fig. 3b and 3e), indicating that the TM segment of cyt b_5_ is required for the formation of the binary complex. The stable binary complex of SCD1-cyt b_5_ is capable of receiving electron from b_5_R, as indicated by the decrease of 340 nm absorbance from NADH (Fig. 3c).

**Fig. 3:**
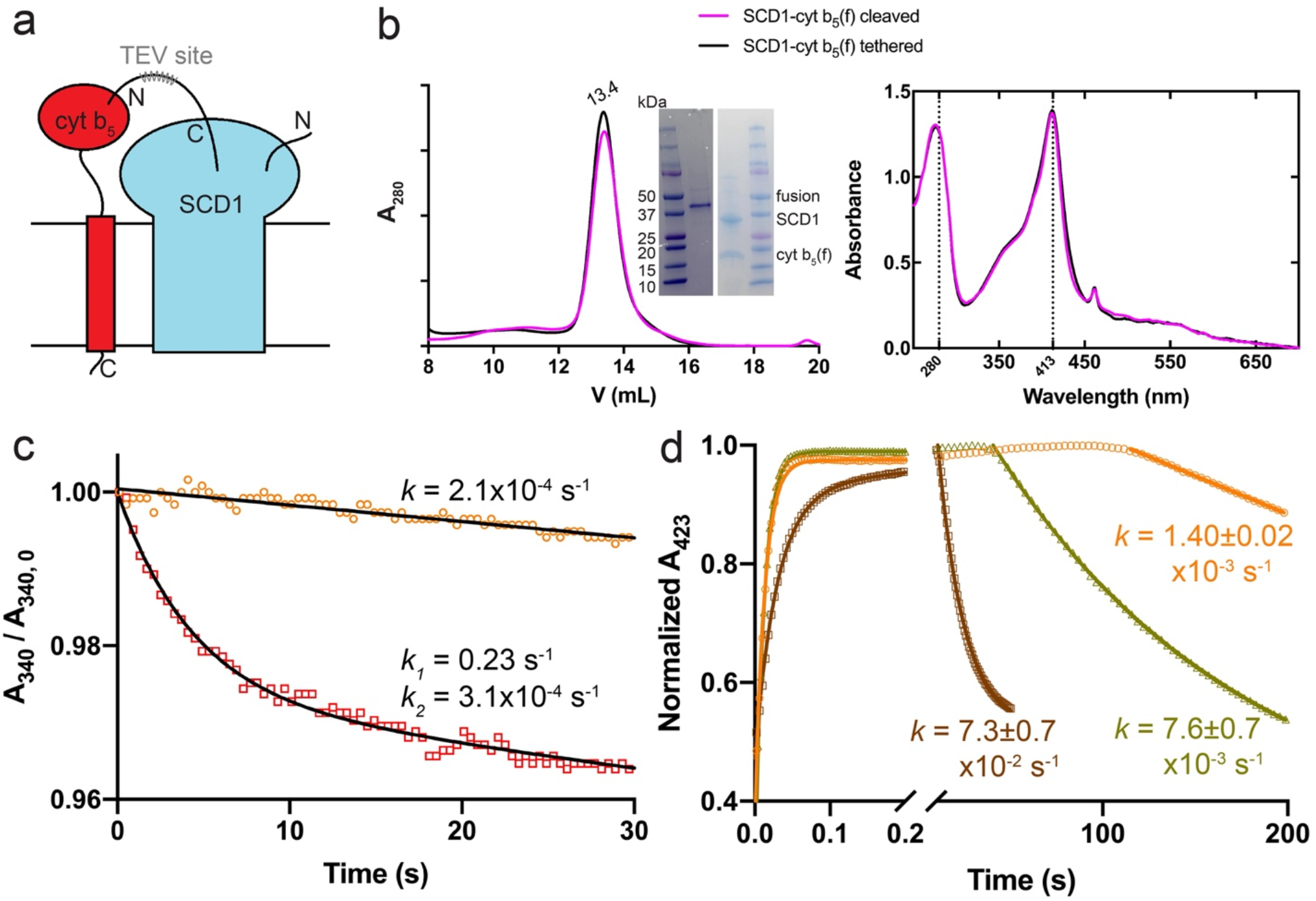
SCD1 and full-length cyt b_5_ can form a stable complex with faster electron transfer kinetics. **a**, Schematic diagram of SCD1-cyt b_5_ fusion constructs. **b**, SEC profile (left), SDS-PAGE image (inset), and UV-Vis spectra (right) of the linker-cleaved (magenta) and tethered (black) fusion of SCD1-cyt b_5_. Almost identical SEC profile and optical spectra suggest the stable assembly between SCD1 and cyt b_5_. **c**, Biphasic kinetics of NADH consumption by b_5_R with SCD1-cyt b_5_ fusion in the presence of substrate stearoyl-CoA (red) compared to the slow linear decrease of NADH in the absence of substrate (orange). The absorbance of NADH at 340 nm (A_340_) was measured and the y-axis is the normalized A_340_ against the initial values (A_340, 0_). Excess NADH was used in the measurements. **d**, Accelerated electron transfer between reduced cyt b_5_ and SCD1 in the SCD1-cyt b_5_ complex (brown) compared to that between the individual cyt b_5_ and SCD1 (yellow) and auto-oxidation of cyt b_5_ (orange). The Soret absorbance of reduced cyt b_5_ at 423 nm was monitored. One molar equivalent of NADH was added, which resulted in the initial rising phases of the fast electron transfer to cyt b_5_ via b_5_R. Rate constants (*k*) are denoted as mean ± SEM calculated from three independent repeats.

We then measured electron transfer rate between the cyt b_5_ and SCD1 (Method and Fig. 3d) by following the optical change of cyt b_5_. The reduced form of cyt b_5_ was followed by its Soret peak at 423 nm. Under anaerobic condition, the first fast-rising phases (t < 20 s) correspond to the reduction of cyt b_5_ by b_5_R after equimolar of NADH was added. After the exhaustion of NADH, the phases of re-oxidation (t > 20 s) of cyt b_5_ by SCD1 appeared. The faster electron transfer rate between cyt b_5_ and SCD1 in the binary complex as opposed to the mixture of individual cyt b_5_ and SCD1 indicates that the binary complex is fully functional and that the formation of the binary complex likely facilitates the alignment and interactions of the two proteins inducive to electron transfer.

### Stable ternary complex between b_5_R, cyt b_5_, and SCD1

Encouraged by the biochemical isolation of the two binary complexes, we examined the production of a ternary complex of SCD1, cyt b_5_ and b_5_R. We took a similar strategy of connecting all three full-length proteins with TEV protease cleavable linkers as shown in Fig. 4a. The fusion protein can be expressed and purified, and eluted as a single peak from a size-exclusion column (Fig. 4b). When the linkers were cleaved, all three proteins stayed together as a ternary complex (Fig. 4b).

**Fig. 4:**
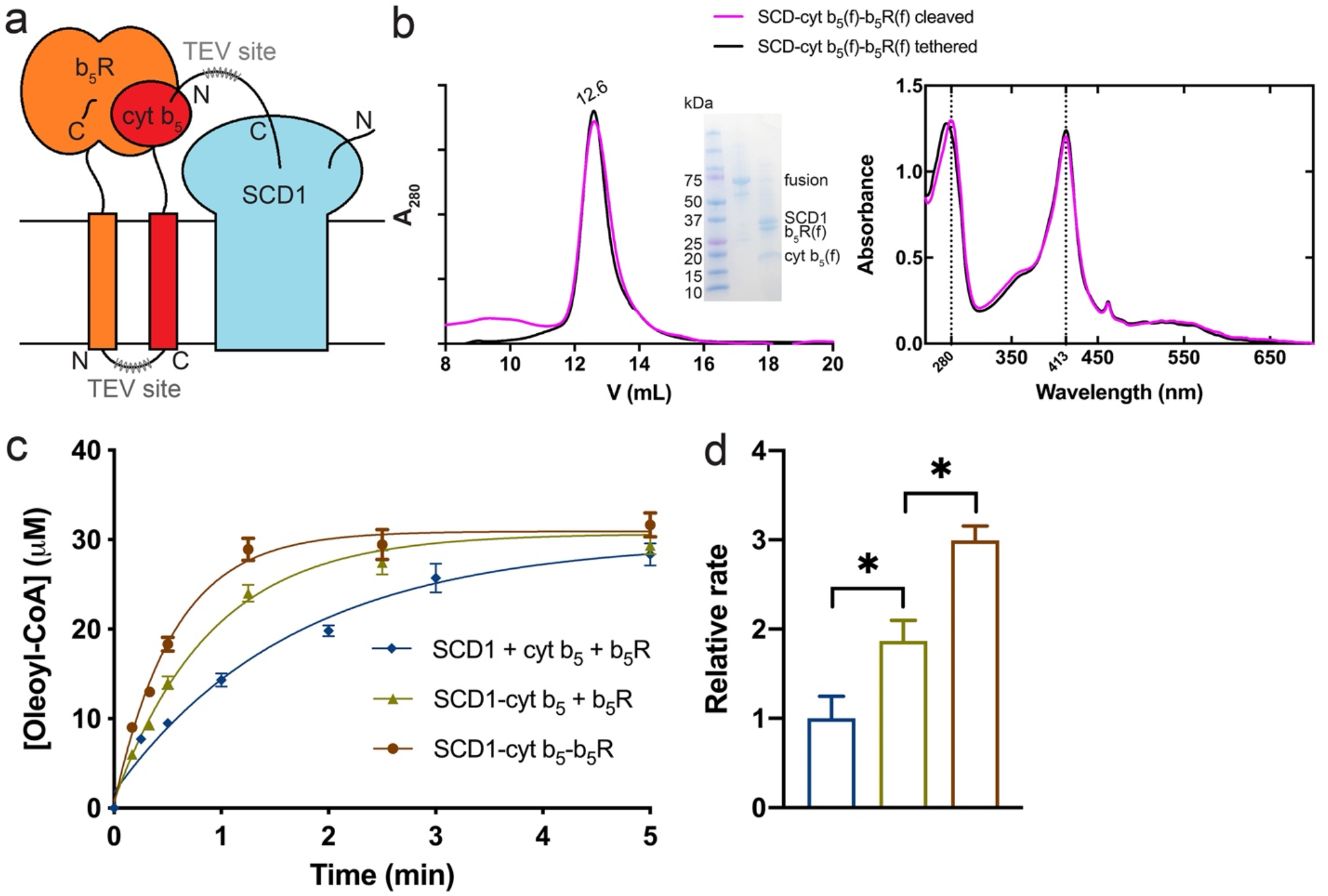
SCD1, full-length cyt b_5_, and full-length b_5_R can form a stable complex with faster electron transfer kinetics. **a**, Schematic diagram of SCD1-cyt b_5_ fusion constructs. **b**, SEC profile (left), SDS-PAGE image (inset), and UV-Vis spectra (right) of the linker-cleaved (magenta) and tethered (black) fusion of SCD1-cyt b_5_-b_5_R. Almost identical SEC profile and optical spectra suggest the stable assembly among SCD1, cyt b_5_, and b_5_R. **c**, Time courses of oleoyl-CoA production in: individual SCD1, cyt b_5_ and b_5_R (blue); the binary complex of SCD1-cyt b_5_ and individual b_5_R (yellow); and the ternary complex of SCD1-cyt b_5_-b_5_R. **d**. Initial rate comparison among conditions in **c**. Asterisks indicate significant different (p < 0.05) in pairwise t-tests. Error bars represent SEM from three independent repeats.

The ternary complex was fully functional, as indicated by the production of oleoyl-CoA when supplied with NADH and stearoyl-CoA (Fig. 4c). The rate of oleoyl-CoA production was significantly faster than those in individual proteins or SCD1-cyt b_5_ binary complex (Fig. 4d), indicating that the ternary complex forms an electron transport chain and enhances the alignment of electron donors and acceptors.

### Models of the binary and ternary complexes

To gain insight into the TM association patterns of SCD1, cyt b_5_ and b_5_R, we generated docking models of their TM regions and did mutational studies to validate key residues involved in the interactions. We first made models of the TM regions of cyt b_5_ and b_5_R as predicted by TMHMM server(Krogh, Larsson et al. 2001). The TM helix of cyt b_5_ was then docked to SCD1, and similarly to that of b_5_R. Best models with fewest clashes were selected and energy minimized (Methods). The selected models of SCD1-cyt b_5_ (TM) and cyt b_5_ (TM) -b_5_R (TM) were aligned with respect to the cyt b_5_ (TM). The aligned model yielded a ternary complex where SCD1 and b_5_R (TM) mounted on different sides of the cyt b_5_ (TM) (Extended Data Fig. 5a). This indicates that the cyt b5 (TM) can interact with both b5R (TM) and SCD1 simultaneously to facilitate the formation of the ternary complex, and the formation of binary or ternary complex is independent. We then performed all-atom MD simulation with the ternary complex model embedded in a 1-palmitoyl-2-oleoyl-sn-glycero-3-phosphocholine (POPC) bilayer (Extended Data Fig. 5a). Three independent 200 ns simulations showed small root-mean-square deviations (RMSD) of 6.6 ± 1.3 Å for cyt b_5_ (TM) and 8.3 ± 1.6 Å for b_5_R (TM) (Extended Data Fig. 5b). The largest root-mean-square fluctuations (RMSF) of cyt b_5_ (TM) and b_5_R (TM) were on residues near surfaces of the membrane, while those buried in the lipidic environment had much less fluctuation (Extended Data Fig. 5c). Collectively, our model was sufficiently stable during MD simulation to be recognized as a ternary complex.

The simulation also revealed the flexible nature of the N-terminal residues (40-60) of SCD1, whose average RMSF = 2.5 Å was significantly larger compared to the average RMSF = 0.3 Å of the rest of protein (Extended Data Fig. 5c). This region could dissociate from a groove on the surface of the soluble domain of SCD1 and expose a positively charged interface for potential interactions with the soluble domain of cyt b_5_. This observation is corroborated by the crystal structures of mouse and human SCD1(Bai, McCoy et al. 2015, Wang, Klein et al. 2015, Shen, Wu et al. 2020), whose N-terminal residues assume different conformations.

To examine if the model is accurate, we introduced single point mutations on the TM helices to the SCD1-cyt b_5_-TEV and cyt b_5_-b_5_R-TEV fusion constructs (Fig 5). Small hydrophobic residues were replaced with a bulky Trp, and polar and large hydrophobic residues were replaced with Ala. Tryptophan substitution on TM domains was shown to weaken or disrupt dimerization of homodimeric ClC transporters while maintaining function of the monomer (Robertson, Kolmakova-Partensky et al. 2010, Chadda, Krishnamani et al. 2016). The mutants were purified and digested with TEV protease and examined on a size-exclusion column. By monitoring the absorbance at 413 nm, where the binary complexes and monomeric cyt b_5_ have the same molar extinction coefficient, we were able to determine the ratio of dimer to monomer (Extended Data Fig. 4).

**Fig. 5:**
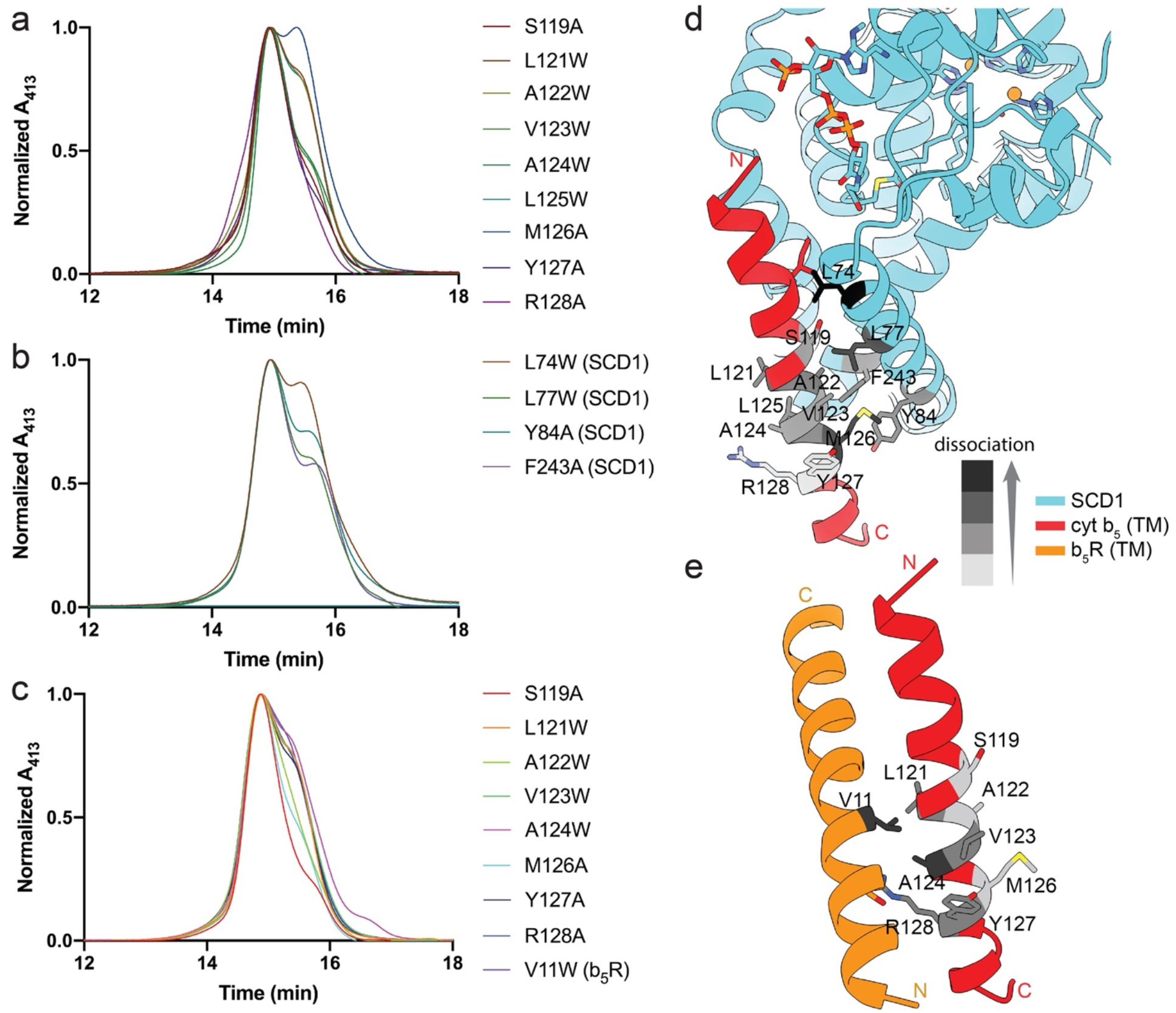
Mutations on the TM domains of SCD1, cyt b_5_, and b_5_R partially disrupt the complex assembly of SCD1-cyt b_5_ and cyt b_5_-b_5_R. SEC profiles of: **a** and **b**, the linker-cleaved SCD1-cyt b_5_ complex; **c**, the linker-cleaved cyt b_5_-b_5_R with mutation on TM domains. Residues on the TM helix of: **a**, cyt b_5_; **b**, SCD1; and **c**, cyt b_5_ or b_5_R were mutated to either alanine from polar or large hydrophobic residues, or bulky tryptophan from small hydrophobic residues. The absorbance at 413 nm from the heme group in cyt b_5_ was monitored in an HPLC system.Docking models of **d**, SCD1 (cyan) and TM helix of cyt b_5_ (red); and **e**, TM helix of cyt b_5_ (red) and b_5_R (orange). The degrees of complex dissociation are evaluated based on profiles in **a**-**c**. Residues important for complex assembly are shown as sidechain stick and are colored per the degree of complex dissociation after mutation. Darker color represents larger effect in disrupting complex assembly.

For the SCD1-cyt b_5_ complex, the M126A on the TM of cyt b_5_ achieved ∼50% of dissociation of cyt b_5_ from SCD1 (Fig. 5a). This is consistent with the docking model in which the M126 interacts with Y84 and F243 on the TMs of SCD1 (Fig. 5d). Introduction of the bulky Trp in the middle of TM helix of cyt b_5_ also destabilized the complex. The strongest effect of Trp mutant was observed on the L121, A122, and V123 of cyt b_5_ (Fig. 5a), which were predicted to be on the side facing SCD1 and involved in the hydrophobic interactions. When the corresponding residues on the SCD1 were mutated, partial dissociation of the complex was also observed (Fig. 5b).

We also mutated residues predicted to be involved in the TM interactions in the cyt b_5_- b_5_R complex. An Arg (R128) on the TM helix of cyt b_5_, which is not energetically favorable to be in a lipidic environment was predicted to interact with a Ser and His on the TM of b_5_R (Fig. 5e), partially neutralizing its charge. The SEC result on the R128A mutant showed that ∼40% of cyt b_5_ fell apart from b_5_R (Fig. 5c). Similar to the results in SCD1-cyt b_5_, Trp mutations to hydrophobic residues (L121, V123, and A124) on TM helix of cyt b_5_ reduced the amount of stable complex (Fig. 5c) by disruption of potential hydrophobic stacking patterns with b_5_R.

Soluble domains of SCD1, cyt b_5_, and b_5_R have very clear complementary electrostatic charge distributions (Extended Data Fig. 6a). Both SCD1 and b_5_R have positively charged surfaces while cyt b_5_ has a negatively charged surface, suggesting that cyt b_5_ may interact one at a time with either b_5_R or SCD1.

We docked the cyt b_5_ separately with SCD1 and b_5_R. Top-ranked docking models brought the heme in cyt b_5_ close to the FAD in b_5_R (Extended Data Fig. 6b) and the diiron center in SCD1 (Extended Data Fig. 6c), which implies direct electron tunneling(Winkler and Gray 2014, Gilbert Gatty, Kahnt et al. 2015) between these cofactors. Based on the predictions, cyt b_5_ engages b_5_R and SCD1 with a similar surface region. To further test this interaction model, we mutated two charged residues on the interface of cyt b_5_ (Extended Data Fig. 6b and 6c) and measured their binding affinities with either b_5_R or SCD1. The affinities to both b_5_R and SCD1 deceased modestly in these mutants (Extended Data Fig. 7), suggesting that these residues are involved in binding with both b_5_R and SCD1.

## Discussion

In summary, we demonstrated that SCD1, cyt b_5_, and b_5_R form a stable ternary complex and that cyt b_5_ forms a stable binary complex with either b_5_R or SCD1. The stable complexes are mediated by TM domains of the proteins, and that the rates of electron transfer greatly increase in these complexes. The formation of stable complexes also suggests that the redox pair of b_5_R and cyt b_5_ may exist as a complex to interact with other downstream proteins.

These results led us to propose a working model of an electron transport chain shown in Fig. 6. Cyt b_5_ and b_5_R form a stable binary complex which further interacts with their target membrane-bound oxidoreductases to form a ternary complex. While the assembly of the binary or ternary complexes is mainly mediated by the interactions between their transmembrane domains, electrostatic interactions between the soluble domains help to steer and further align the donor-acceptor redox pairs. Cyt b_5_, while its single TM is sandwiched between b_5_R and SCD1 to stabilize the ternary complex, its soluble domain is mobile and can interact alternatively between b_5_R and SCD1 to relay electrons. A recent study revealed that lipid solvation energies drive the association and dissociation of a dimeric ClC transporter (Chadda, Bernhardt et al. 2021), and we surmise that the formation of the ternary SCD1-cyt b5-b5R complex maybe influenced by the lipid content of the ER membrane.

**Fig. 6:**
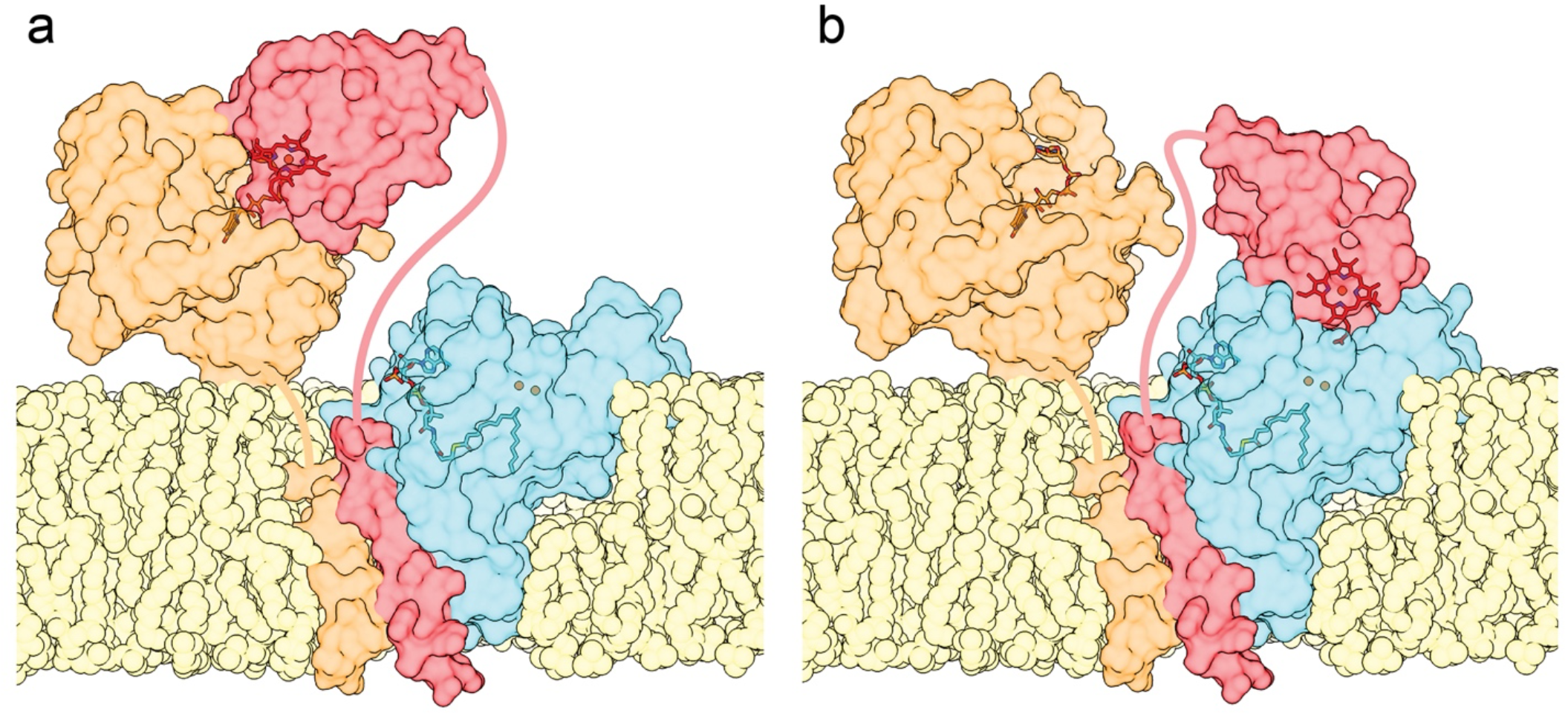
Model of SCD1-cyt b_5_-b_5_R complex. SCD1 (cyan), cyt b_5_ (red), and b_5_R (orange) are shown as translucent surface highlighting diiron center and acyl-CoA in SCD1, heme in cyt b_5_ (deep red), and FAD in b_5_R (brown). SCD1 form complex with the TM helices of cyt b_5_ and b_5_R in lipid bilayer (yellow). A flexible linker connecting the soluble domain of cyt b_5_ and b_5_R to their TM helices allows the transition of two states: **a**, cyt b_5_ receiving an electron from b_5_R; and **b**, cyt b_5_ delivering an electron to SCD1.

The association of the soluble domains of b_5_R and cyt b_5_, which harbor the cofactors for electron transport, is weak (*K*_*D*_ ∼ µM, Extended Data Fig. 7) and likely transient. Anchoring them to the membrane would constrain their diffusion to the 2-dimensional lipid bilayer removing the third dimension of free diffusion in cytosol and thus enhances their ability to form the binary complex for electron transfer. Our results indicate that the interactions between the b_5_R, cyt b_5_, and SCD1 transmembrane domains greatly enhance the stability of the binary and ternary complexes. Such complexes bring the soluble domains of SCD1, cyt b_5,_ and b_5_R in proximity and may rigidify each component for the optimal alignment required for efficient electron transfer.

Soluble domains b_5_R and cyt b_5_ are connected to their TM helix with a long linker predicted to be unstructured. This may afford certain degrees of conformational freedom allowing the soluble domains to contact with one another for efficient electron transfer. Such flexibility may also allow cyt b_5_ to shuttle electrons between b_5_R and SCD1 in the ternary complex.

Recent studies(Ahuja, Jahr et al. 2013, Yamamoto, Dürr et al. 2013, Zhang, Huang et al. 2015, Jeřábek, Florián et al. 2016, Zhang, Huang et al. 2016, Yamamoto, Caporini et al. 2017, Yamamoto, Caporini et al. 2017) using nuclear magnetic resonance and molecular dynamic (MD) simulation have shown interactions between TM helix of cyt b_5_ and that of cyt P450, an oxidoreductase with a single TM helix. The authors found that when both full-length cyt b_5_ and cyt P450 were incorporated into a single lipid nanodisc(Zhang, Huang et al. 2016), their TMs could interact and that the interactions facilitate electron transport between the soluble domains of cyt b_5_ and cyt P450(Zhang, Huang et al. 2015). Interestingly, a conserved motif on cyt b_5_ (L121 – L125) that was thought to interact with cyt P450 in nanodiscs(Yamamoto, Caporini et al. 2017) is also identified in the current study to mediate interactions with SCD1 and b_5_R.

Understanding the interactions of SCD1, cyt b_5_, and b_5_R and ultimately obtaining structures of the ternary complex in different redox states will help our understanding of SCD1 and other oxidoreductases that rely on cyt b_5_ and b_5_R. The knowledge also provides insights into designing small molecules that target the interactions between the transmembrane domains which have not been the target regions of the proteins.

## Methods

### DNA constructs

The cDNA of mouse SCD1, full-length cyt b_5_, and full-length b_5_R were codon optimized and synthesized. Fusions of SCD1-cyt b_5_, cyt b_5_-b_5_R, SCD1-cyt b_5_-b_5_R were generated by PCR. The linkers between each domain were either a TEV protease site (ENLYFQ/G) for the cleavable fusion or a flexible linker (GGSGGGSG) for the non-cleavable fusion. The SCD1 and SCD1-cyt b_5_ fusions were cloned into a pEG BacMam vector with a TEV protease site prepended to a C-terminal GFP tag. Because the C-terminus of b_5_R ends on the interface of the FAD-binding domain and NADH-binding domain, no extra residue should be introduced after the C-terminus of b_5_R domain to preserve its functional integrity. Therefore, SCD1-cyt b_5_-b_5_R was cloned into a pEG BacMam vector with an N-terminal GFP tag appended with a TEV protease site. For immunofluorescence imaging and coimmunoprecipitation assays, cyt b_5_ and cyt b_5_-b_5_R with a N-terminal Myc tag, and b_5_R with a N-terminal HA tag were cloned into a pEG BacMam vector. For large-scale protein expression and purification, cyt b_5_, b_5_R, and cyt b_5_-b_5_R were cloned into a pFastBac Dual vector with an octa-histidine tag and TEV protease site. For Octet binding assays, the soluble domains of cyt b_5_ (4 - 89) and b_5_R (24 - 301) were cloned into a pET vector.

### Immunofluorescence imaging

The HEK 293S cells in *FreeStyle 293* media (Invitrogen/Thermo Fisher) supplemented with 2% fetal bovine serum (FBS; Sigma) were plated one day before transfection onto glass coverslips coated with poly-lysine in a 24-well plate. The cDNAs in pEG BacMam vectors were transfected into cells with Lipofectamine 2000 (Invitrogen/Thermo Fisher) per manufacturer’s instructions. About 24 h after transfection, cells were fixed with 2% paraformaldehyde for 10 min and then washed three times with phosphate buffered saline (PBS). Cells were blocked and permeabilized with PBSAT (PBS + 1% bovine serum albumin, BSA + 0.1% Triton X-100) for 10 min. For cells expressing only GFP-tagged proteins, coverslips were washed three times with PBS and mounted onto glass slides with *ProLong* Diamond (Invitrogen/Thermo Fisher). For cells expressing Myc and/or HA tagged proteins, primary antibodies against Myc and/or HA tag diluted in PBSAT were added and incubated for 45 min at room temperature (RT). Coverslips were washed three time with PBS before the incubation with Alexa Fluor 555 (for Myc tag) and/or Alexa Fluor 647 (for HA tag) conjugated secondary antibodies (Invitrogen/Thermo Fisher) diluted in PBSAT for 30 min at RT. Finally, coverslips were washed and mounted as mentioned before.

Confocal images were acquired with a Zeiss LSM-710 confocal microscope using a 63× oil immersion objective (Zeiss, Plan-Apochromat 63×/1.4 Oil DIC M27) with *Immersol* 518F immersion oil (Zeiss). Alexa Fluor 647, Alexa Fluor 555, and GFP were detected sequentially with 633 nm HeNe laser, 561 nm diode-pumped solid-state laser, and 488 nm Argon laser. Crosstalk between the channels was avoided by adjusting emission regions. Single optical sections at a resolution of 1024×1024 pixels were acquired at two different zoom levels (1.5× and 4×).

### Coimmunoprecipitation

The HEK 293S cells were plated one day before transfection in a 6-well plate. The pEG BacMam vectors containing target cDNAs were transfected with Lipofectamine 2000 (Invitrogen/Thermo Fisher) and incubated for 2 days. Cells on the plate were washed in PBS before scraping. Cell membranes were solubilized in lysis buffer (20 mM HEPES, pH 7.5, 150 mM NaCl, 10% glycerol) plus 0.2% Triton X-100 and Protease Inhibitor Cocktail (Roche) for 1 h at 4°C. Cell debris were pelleted by centrifugation. The supernatants of cell lysate were incubated with either pre-equilibrated GFP nanobody-conjugated NHS-Activated Sepharose 4 Fast Flow Agarose (GE Healthcare) or *Pierce* Protein A Agarose (Invitrogen/Thermo Fisher) with rabbit anti-Myc antibodies for 30 min at 4°C. The resins were extensively washed in lysis buffer plus 0.1% Triton X-100 within 5 min at 4°C. 4× Laemmli Sample Buffer (Bio-Rad) was added and samples were run in SDS-PAGE without extra elution steps. Bands of target proteins were visualized by western blotting with mouse anti-GFP (Invitrogen/Thermo Fisher), anti-Myc, and anti-HA antibodies as primary antibodies and IRDye-800CW anti-mouse IgG (Licor) as secondary antibody. Images were taken on an Odyssey infrared scanner (Licor).

### Large-scale expression and purification of proteins

Expression of SCD1-containing proteins (SCD1, SCD1-cyt b_5_, and SCD1-cyt b_5_-b_5_R) were conducted in HEK 293S cells using the BacMam system(Goehring, Lee et al. 2014). Baculovirus were generated from pEGBacMam vectors with target cDNAs and amplified in Sf9 (*Spodoptera frugiperda*) cells. HEK 293 cells were maintained in *FreeStyle 293* media (Invitrogen/Thermo Fisher) supplemented with 2% FBS (Sigma) in a 37°C incubator with 8% CO_2_ atmosphere at 100 rpm. Baculovirus after three passages (P3) were added to HEK 293S cells at a density of 3 × 10^6^ mL^-1^ at a 7.5% v/v ratio and incubated overnight before adding 10 mM sodium butyrate and lowering temperature to 30°C. Media were supplemented with transferrin and ferric chloride as described previously(Shen, Wu et al. 2020). 0.5 mM δ-aminolevulinic acid and 100 µM riboflavin were added in media to enhance the biosynthesis of heme and FAD group, respectively. Three days after infection, cells were harvested and resuspended in lysis buffer plus Protease Inhibitor Cocktail (Roche), 1 mM phenylmethylsulfonyl fluoride (PMSF), 5 mM MgCl_2_ and DNase I. Solubilization of cell membranes was achieved by incubating with 30 mM n-Dodecyl-β-D-Maltopyranoside (DDM, Anatrace) for 2 h at 4°C under gentle agitation. Target proteins in supernatants were captured by GFP nanobody resins. After washing the resins with 20 CV washing buffer (lysis buffer plus 1 mM DDM), proteins were released by TEV protease digestion during which TEV protease site linkers in the cleavable fusions were also cleaved. Eluents were concentrated (Amicon 50-kDa cutoff, Millipore) and loaded onto a size-exclusion column (Superdex 200 10/300 GL, GE Health Sciences) equilibrated with FPLC buffer (20 mM HEPES, pH 7.5, 150 mM NaCl, 1 mM DDM).

Expression of full-length cyt b_5_, full-length b_5_R, and cyt b_5_-b_5_R was conducted in High Five (*Trichoplusia ni*) cells using the Bac-to-Bac system. Baculovirus were generated from pFastBac Dual vectors with target cDNAs. 1.5% v/v of P3 virus was added to cells at a density of 3 × 10^6^ mL^-1^. The δ-aminolevulinic acid (Santa Cruz) and/or riboflavin (Sigma) were supplemented in media as mentioned before. Cells were harvested three days after infection. Purification procedure was similar to that for HEK 293 cells except for the usage of cobalt-based affinity resin (Talon, Clontech) to capture His-tagged proteins.

### UV-Vis spectroscopy and enzymatic assays

UV-Vis spectra were recorded using a Hewlett-Packard 8453 diode-array spectrophotometer (Palo Alto, CA). The time courses of the NADH consumption at 340 nm and the spectral change of cyt b_5_ heme at 423 nm were obtained with an Applied Photophysics (Leatherhead, UK) model SX-18MV stopped-flow instrument. The observed rates, *k*_obs_, were obtained by fitting the time courses to either 1- or 2-exponential functions.

Continuous turnover reactions of SCD1, the binary complex of SCD1-cyt b_5_, and the ternary complex of SCD1-cyt b_5_-b_5_R were performed similarly as previously described. Briefly, 3 µM of SCD1 plus equimolar of cyt b_5_ and b_5_R, SCD1-cyt b_5_ plus equimolar of b_5_R, and SCD1-cyt b_5_-b_5_R in FPLC buffer were incubated with substrate stearoyl-CoA (Sigma). NADH was added to start reaction. Aliquots of reaction mixtures were retrieved and quenched at different time points, and analyzed in high-performance liquid chromatography (HPLC). The initial rates were calculated by linear fitting of time courses within 1 min after reaction started.

### Modeling of binary and ternary complex

The TM regions of cyt b_5_ and b_5_R were prediction by TMHMM server(Krogh, Larsson et al. 2001) and residue 108 - 134 of cyt b_5_ and residue 1 - 28 of b_5_R were used to model a TM helix a in I-TASSER server(Yang and Zhang 2015). The helical models with highest C-score were chosen for docking in Memdock(Hurwitz and Wolfson 2021). The diiron-containing mouse SCD1 structure (PDB ID: 6WF2) was used for docking. The resulting top 5 models with smallest Memscore were manually assessed based on topology and orientation in membrane predicted by PPM server(Lomize, Pogozheva et al. 2012). The selected models of SCD1-cyt b_5_ (TM) and cyt b_5_ (TM)-b_5_R (TM) were placed in a membrane bilayer of POPC and energy-minimized and equilibrated in the CHARMM36 force field(Huang, Rauscher et al. 2017). The two binary models were aligned to cyt b_5_ (TM) to generate a ternary model.

We used SWISS-MODEL server(Waterhouse, Bertoni et al. 2018) to construct homology models of mouse soluble cyt b_5_ (1 - 89) and b_5_R (24 - 301). The surface electrostatic potential of SCD1, cyt b_5_, and b_5_R were calculated in APBS(Jurrus, Engel et al. 2018). Dockings of soluble cyt b_5_ and b_5_R, and SCD1 and soluble cyt b_5_ were performed in HADDOCK server(van Zundert, Rodrigues et al. 2016) using a flexible docking protocol. Charged residues on the presumed binding interfaces were defined as active residues to restrain their proximity during docking process. The FAD in b_5_R and heme in cyt b_5_ were also included as part of the interfaces. All structure figures were prepared in Pymol (Schrödinger LLC.) or ChimeraX(Pettersen, Goddard et al. 2021).

### Size-exclusion chromatography of mutants

Mutations on the TM domains of SCD1, cyt b_5_, and b_5_R were introduced to the constructs of linker-cleavable fusions of SCD1-cyt b and cyt b_5_-b_5_R by QuikChange site-directed mutagenesis. All mutations were confirmed by sequencing. Expression and purification were done similarly to the wild-type (WT) fusions. Purified proteins were loaded onto a SEC column (SRE-10C SEC-300, Sepax) in an HPLC system with a diode-array detector (SPD-M20A, Shimadzu). Samples were run in FPLC buffer at a flow rate of 0.75 mL/min and monitored at 423 nm. Elution profiles were normalized to the peak corresponding to either SCD1-cyt b_5_ or cyt b_5_-b_5_R complex for comparison.

### Molecular dynamic simulation

The simulation system was prepared in CHARMM-GUI Membrane Builder(Jo, Kim et al. 2007, Jo, Kim et al. 2008, Jo, Lim et al. 2009, Wu, Cheng et al. 2014, Lee, Patel et al. 2019) with the ternary model of SCD1-cyt b_5_(TM)-b_5_R(TM). The structure was protonated at neutral pH and was embedded in a POPC bilayer. The system was solvated with TIP3P water and 150 mM NaCl (including neutralization ions). The final system had a size of 100 Å × 100 Å × 107 Å.

The simulations were performed in Gromacs v.2021.2(Abraham, Murtola et al. 2015) with the CHARMM36 force field(Lee, Cheng et al. 2016) in the isothermal-isobaric (NPT) ensemble. The system temperature was maintained at 303.15 K using the Langevin temperature coupling method with a friction coefficient of 1 ps^-1^. The semi-isotropic Nosé-Hoover Langevin-piston method was used to maintain the pressure at 1 atm. The 10–12 Å force-based switching was used for the Lennard-Jones interaction. The particle mesh Ewald method was used for the long-range electrostatic interactions. Three 200 ns unrestrained production simulations were performed using a timestep of 2 fs. Trajectory data were analyzed with the utilities in Gromacs(Abraham, Murtola et al. 2015).

### Octet Biolayer Interferometry

Soluble cyt b_5_ and b_5_R were expressed in BL21(DE3) cells with pET vectors containing target cDNA. Mutations in cyt b_5_ were introduced by QuikChange site-directed mutagenesis and were confirmed by sequencing. Protocols for expression of these proteins were adapted from previous reports(Mulrooney and Waskell 2000, Bando, Takano et al. 2004). Purification procedure was similar to that for full-length cyt b_5_ and b_5_R as mentioned before.

Biolayer Interferometry (BLI) assays were performed at 30 °C under constant shaking at 1000 rpm using an Octet system (FortéBio). The immobilization of ligand proteins on amine reactive second-generation (AR2G) biosensors (Sartorius) was done following manufacturer’s instruction. Briefly, biosensor tips were activated in 20 mM 1-ethyl-3-[3-dimethylaminopropyl]carbodiimide hydrochloride (EDC) and 10 mM N-hydroxysulfosuccinimide (Sulfo-NHS) for 300 s. Then the tips were loaded with soluble cyt b_5_ at a concentration of 5 µg/mL in FPLC buffer for 600 s. The tips were quenched in FPLC buffer plus 1 M ethanolamine for 300 s. The tips with immobilized ligands were equilibrated in FPLC buffer plus 0.1% BSA to reduce non-specific binding. Then, they were transferred to wells of a concentration gradient (5, 2.5, and 1.25 µM) of analysts (soluble b_5_R or SCD1) in buffer B for association and returned to the equilibration wells for dissociation. Binding curves were aligned and corrected with the channel of no analyst protein. The association and disassociation phases were fitted with 1-exponential function to extract *k*_*on*_ and *k*_*off*_ of the binding, which were used to calculate dissociation constant *K*_D_.

## Acknowledgments

We thank Dr. Theodore G Wensel for the access of Zeiss confocal microscope and western blot-related materials; Dr. Melina A Agosto for the help with confocal microscopy and co-IP experiments; and Joshua I Rosario Sepulveda for the help with western blot. This work was supported by grants from NIH (DK122784 to M.Z. and A.T., HL086392 and GM098878 to M.Z.), and Cancer Prevention and Research Institute of Texas (R1223 to M.Z.).

## Author Contributions

M.Z., A.T., J.S., and G.W. conceived the project. J.S. and G.W. conducted experiments. J.S. and M.Z. wrote the initial draft and all authors participated in revising the manuscript.

## Corresponding authors

Ming Zhou, Ah-Lim Tsai

---

**Scheme 1:**
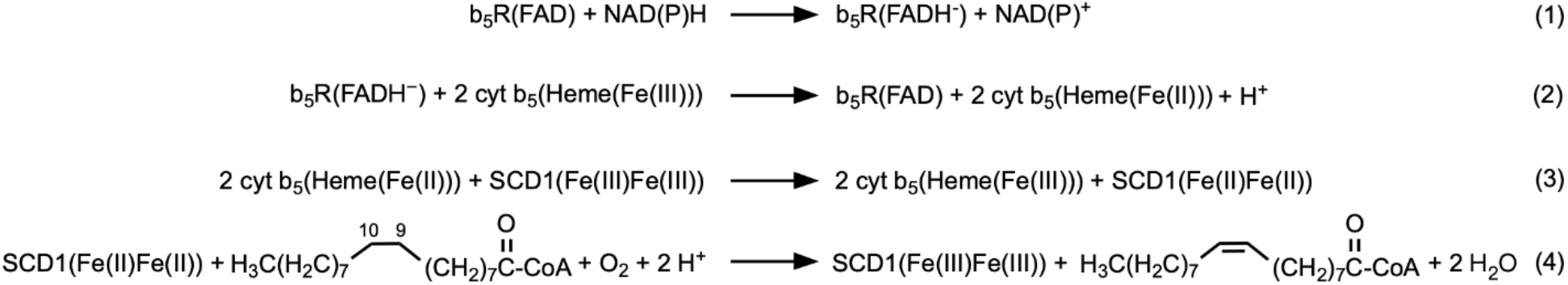
Stepwise reactions in the electron transport chain of b_5_R, cyt b_5,_ and SCD1.

## Extended Data

**Extended Data Fig. 1:**
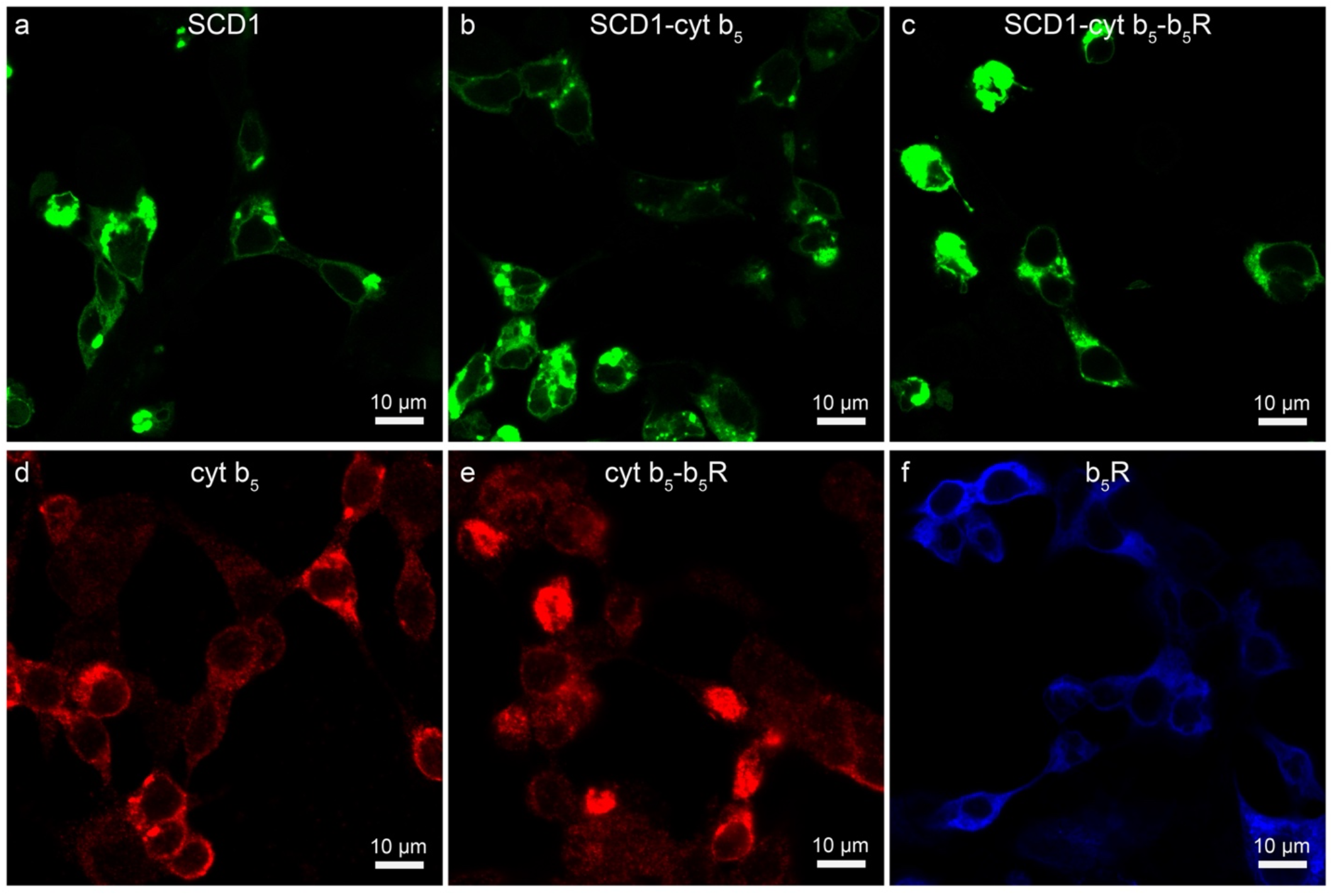
Expression and subcellular distribution of SCD1, cyt b_5_, b_5_R, and their binary and ternary fusions. Confocal microscopy image of HEK293 cells expressing **a**, SCD1-GFP; **b**, SCD1-cyt b_5_-GFP; **c**, GFP-SCD1-cyt b_5_-b_5_R; **d**, Myc-cyt b_5_; **e**, Myc-cyt b_5_-b_5_R; and **f**, HA-b_5_R.

**Extended Data Fig. 2:**
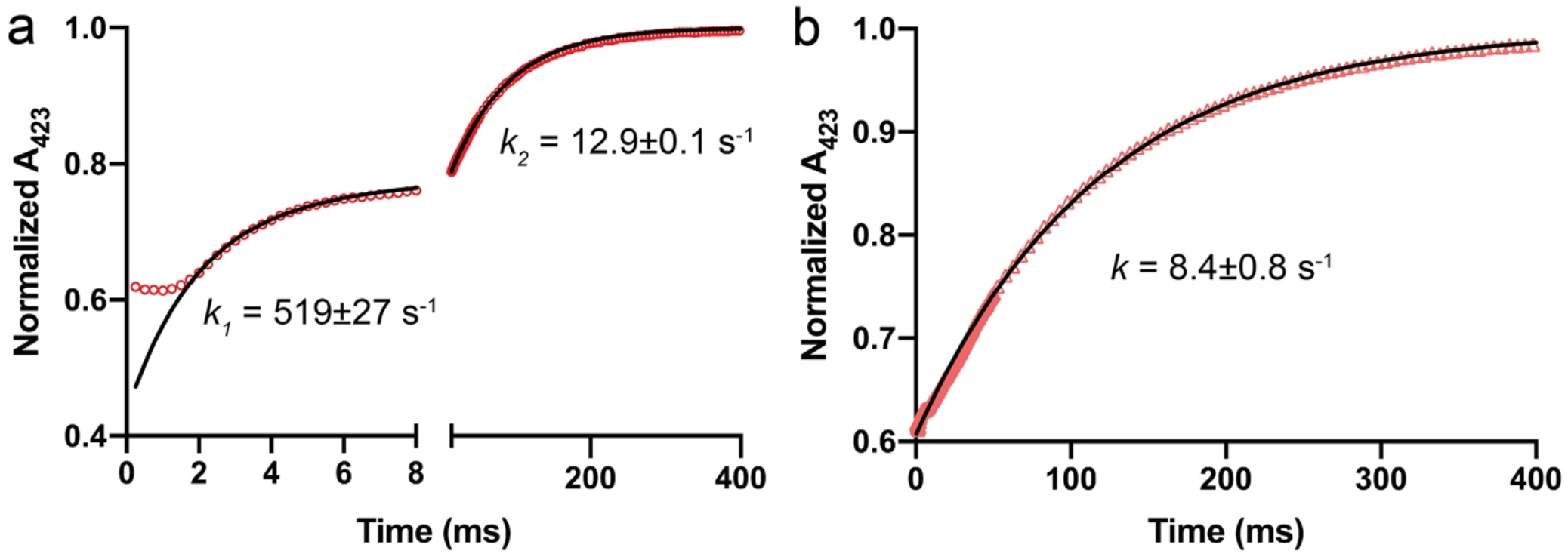
Representative time courses of electron transfer between b_5_R and cyt b_5_ in **a**, cyt b_5_-b_5_R complex; and **b**, individual cyt b_5_ and b_5_R. The reduction of cyt b_5_ was followed at 423 nm. The time course is biphasic for cyt b_5_-b_5_R complex with a fast phase (*k*_*1*_) and a slow phase (*k*_*2*_), while monophasic (*k*) for the individual cyt b_5_ and b_5_R. Red dots represent experimental data and black curves the 1-exponential fit. Rate constants are denoted as mean ± SEM of three independent repeats.

**Extended Data Fig. 3:**
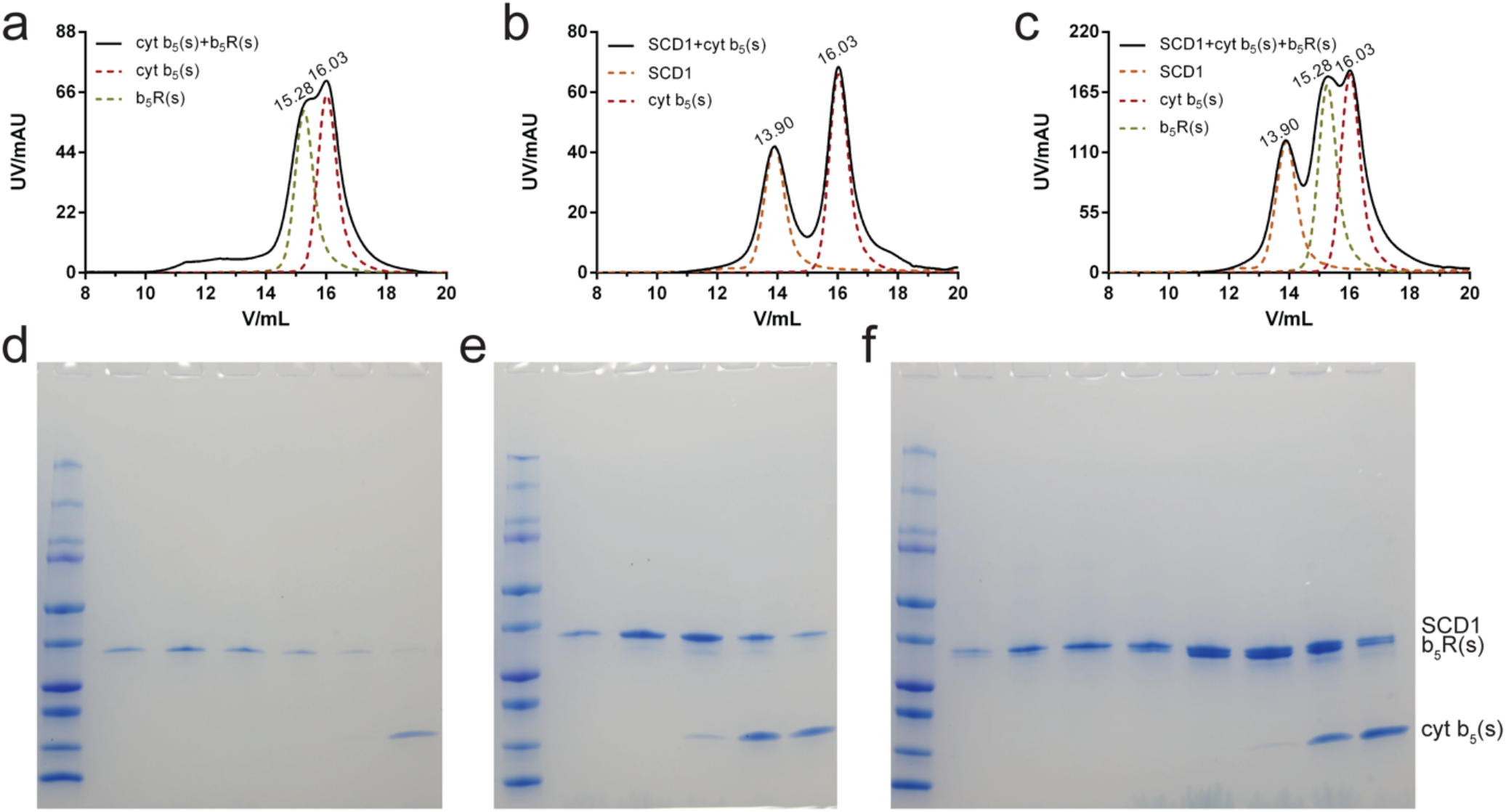
SCD1, soluble cyt b_5_, and b_5_R cannot form stable complex in SEC. **a** and **d**, SEC profile of a mixture of soluble cyt b_5_ and b_5_R, and SDS-PAGE of fractions within the elution volume of 14-17 mL. **b** and **e**, SEC profile of a mixture of SCD1 and soluble cyt b_5_, and SDS-PAGE of fractions within the elution volume of 14-17 mL. **c** and **f**, SEC profile of a mixture of SCD1, soluble cyt b_5_ and b_5_R, and SDS-PAGE of fractions within the elution volume of 13-17 mL.

**Extended Data Fig. 4:**
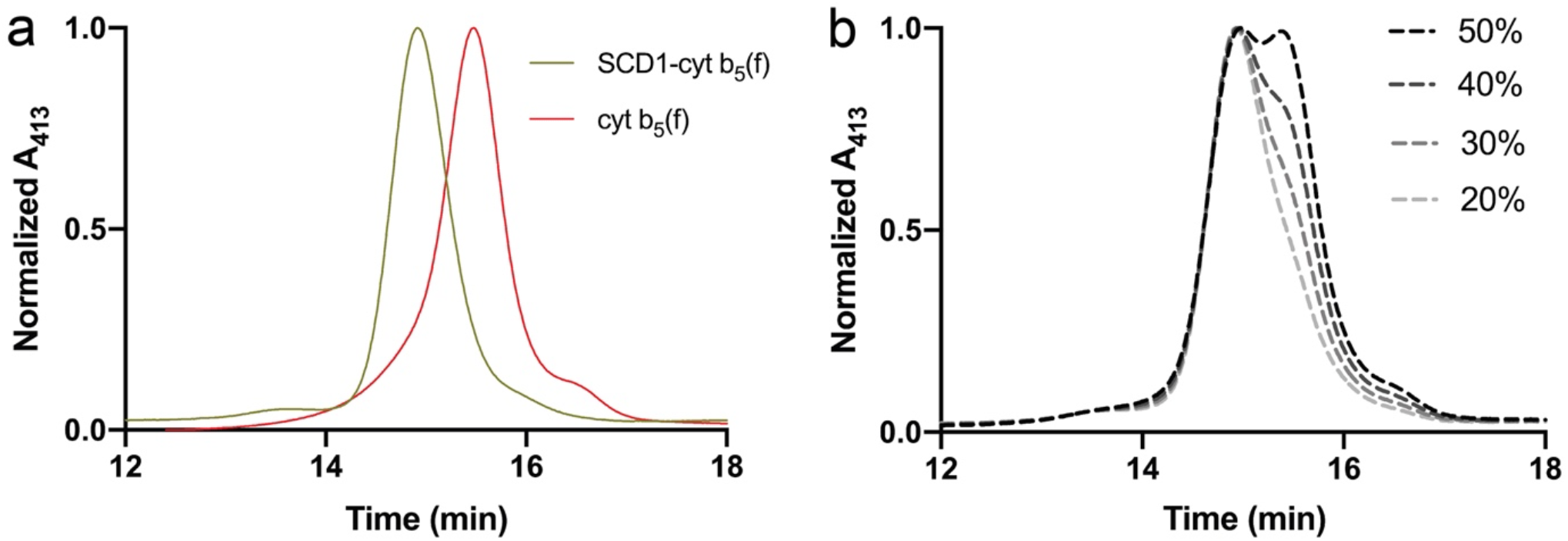
**a**, SEC profiles of SCD1-cyt b_5_ (yellow) complex and full-length cyt b_5_ (red) in an HPLC system monitoring absorbance at 413 nm. **b**, Simulated profiles of partially dissociated complex based on profiles in **a**. Percentages represent the molar ratio of dissociated cyt b_5_ to all heme-containing proteins, including SCD1-cyt b_5_ complex and individual cyt b_5_.

**Extended Data Fig. 5:**
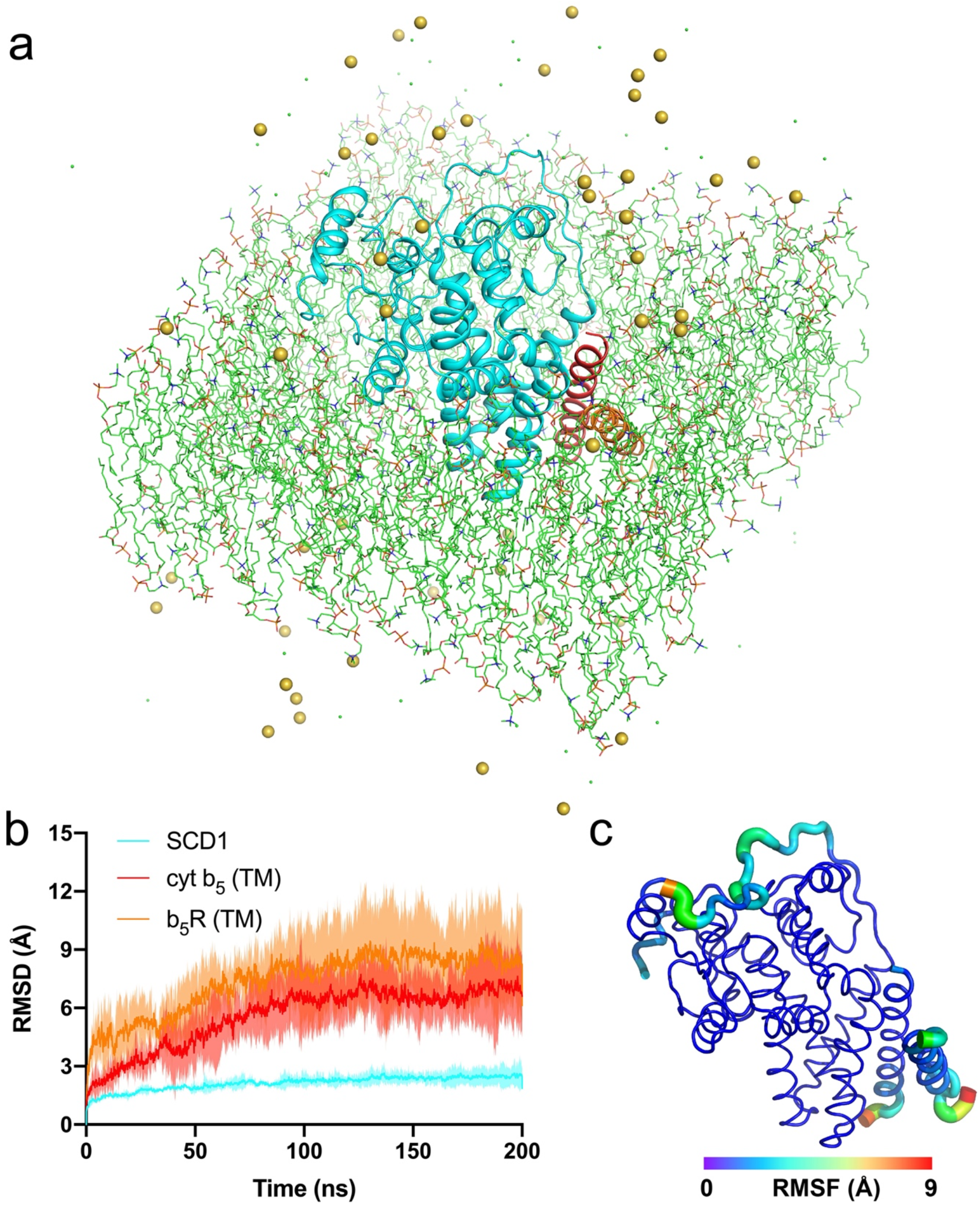
MD simulation of the SCD1-cyt b_5_-b_5_R complex. **a**, The environment for the simulation of the complex. The docking model of the ternary complex (shown as cartoon; SCD1 in cyan, the TM helix of cyt b_5_ in red, and the TM helix of b_5_R in orange) are embedded in a lipid bilayer of POPC (shown as line). The solvent contains 150 mM NaCl. Na^+^ and Cl^-^ were shown as yellow and green spheres, respectively. Water molecules are hidden for clarity. **b**, The time courses of root-mean-square deviation (RMSD) of the backbone atoms in SCD1 (cyan), the TM helix of cyt b_5_ (red), and TM helix of b_5_R (orange) of the complex. For the calculation of RMSD, the backbone atoms of SCD1 were aligned to the initial complex structure. The time courses are shown as solid line (mean) with shaded region (standard deviation) calculated from three independent runs. **c**, The root-mean-square fluctuation (RMSF) of backbone C_α_ atoms of the complex. The backbones are displayed as tubes and colored in a rainbow spectrum as per their RMSF values. The diameter of the tubes also correlates to the RMSF values. The RMSF values are averages calculated from three 200 ns simulations.

**Extended Data Fig. 6:**
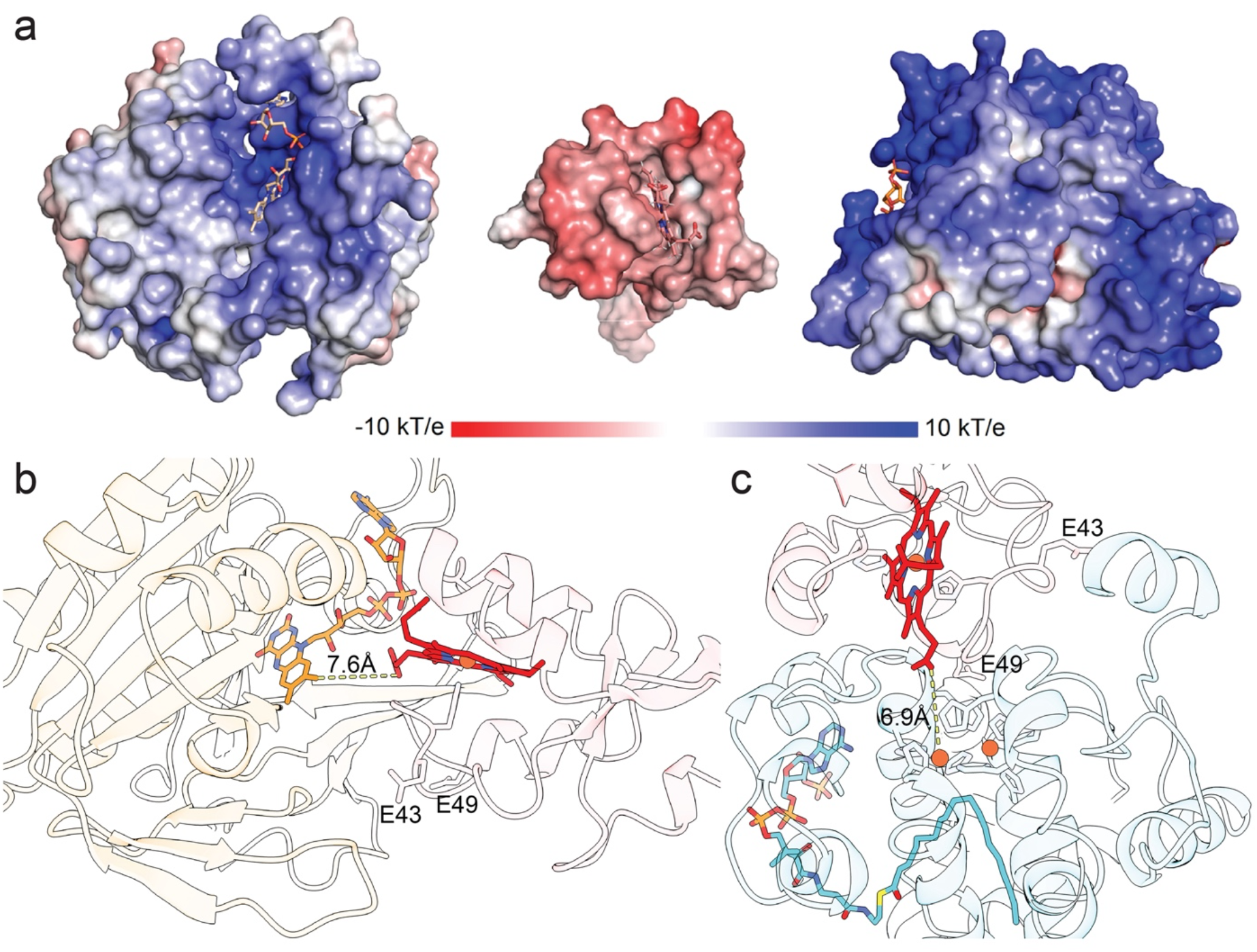
**a**, Electrostatic potential of soluble domain of b_5_R (left), cyt b_5_ (middle), and SCD1 (right). The binding interfaces are shown as surface colored from red to blue to represent potential from −10 kT/e to +10 kT/e. Prosthetic group and ligand are shown as stick. Docking models of: **b**, the soluble domains of cyt b_5_ (red) and b_5_R (orange); and **c**, SCD1 (cyan) and the soluble domain cyt b_5_ (red). The FAD in b_5_R, heme in cyt b_5_, and diiron center and acyl-CoA in SCD1 are highlighted. The closest distances between redox centers in the models are labeled. Two residues (E43 and E49) of cyt b_5_ on the binding interfaces in both cyt b_5_-b_5_R and SCD1-cyt b_5_ complex are shown as sidechain sticks.

**Extended Data Fig. 7:**
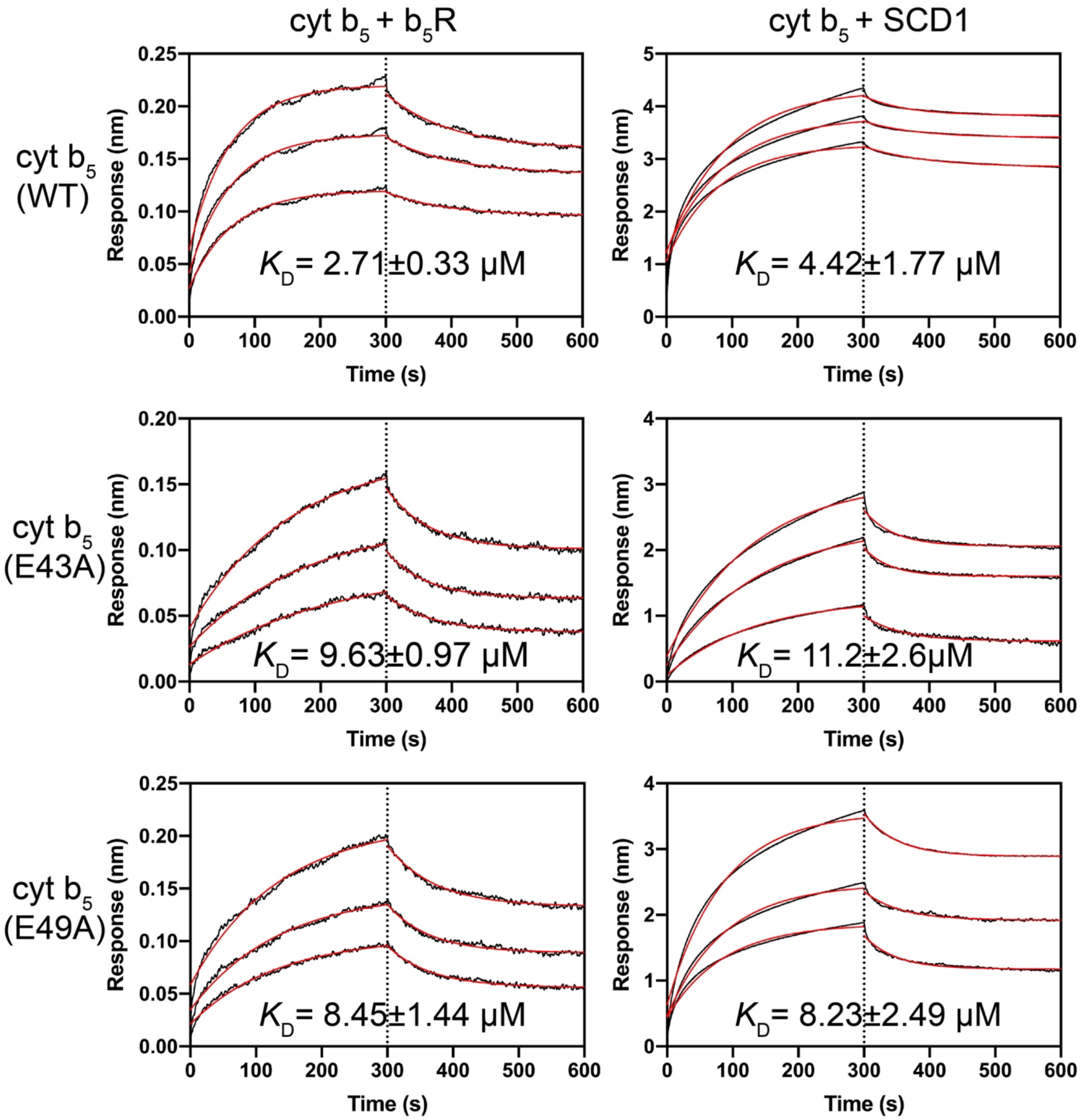
Binding of wild-type (WT) and surface mutants (E43A and E49A) of cyt b_5_ to b_5_R (left panel) or SCD1 (right panel) measured in Octet BLI. *K*_D_ values were calculated as described in Methods.

